# Accurate interdomain contacts in mixed folded proteins from NMR-guided coarse-grained simulations

**DOI:** 10.1101/2025.11.21.689726

**Authors:** Billy Hobbs, Noor Limmer, Gwendolyn L. Clenshaw, Felipe Ossa, Theodoros K. Karamanos

## Abstract

Intrinsically disordered, low-complexity regions frequently cooperate with folded domains to mediate protein–protein interactions, yet accurately describing these mixed folded–disordered systems remains challenging. To visualize these mixed folded proteins, experimentally guided coarse-grained (CG) molecular dynamics simulations are often employed to extend the timescales required to capture the complex dynamics in play. However, the minimalistic nature of these approaches often compromises structural accuracy and can lead to inaccurate inter-domain interactions. Here we introduce backbone dihedral terms directly derived from NMR chemical shift data in CG-simulations to characterize the open state of a mixed-folded construct of the anti-aggregation chaperone DNAJB6 that contains a folded J-domain and a disordered GF linker. By tuning residue-specific backbone dihedral parameters to match NMR-derived secondary-structure propensities of the linker in CG-simulations, we generate conformational ensembles that yield accurate interdomain contact maps. In agreement with analysis of NMR relaxation data, the resulting ensembles show that even in the nominally open state the linker experiences motions that resemble those of the closed state driven by hydrophobic residues in GF. More generally we show that by expanding CG-simulations to allow them to capture both local and global structural properties, physically relevant interdomain contacts can be retrieved.

## Introduction

Low complexity regions (LCRs) in proteins have attracted significant research interest as they serve crucial roles in important biological processes.^1,2^ In many cases intrinsically disordered LCRs mediate interactions between folded domains by increasing the local concentration of the later or by acting as hotspots for protein-protein interactions.^2^ However, and despite their importance, the atomic level characterization of low complexity regions still represents a significant task due to their inherent dynamics and similar spectroscopic signatures that complicate analysis by the majority of structural biology methods.

Computational methods have provided crucial insights into the conformational ensembles of intrinsically disordered regions (IDRs), often guided by experimental data.^3^ However, the millisecond timescales required to adequately sample IDR conformational space constitute all-atom molecular dynamics (MD) simulations prohibitively expensive, motivating the development of coarse-grained (CG) models.^4,5^ While CG simulations successfully capture global ensemble properties such as chain compaction, their minimalistic nature often complicates the incorporation and/or extraction of critical sequence-specific features. In particular, residue-level secondary-structure propensities—known to be functionally important in many IDRs—are difficult to represent accurately, leading to incorrect or non-physical inter-residue and interdomain contact patterns.^6^ As a result, CG approaches frequently struggle to describe the interplay between folded domains and disordered regions.^6^ Here, using a J-domain chaperone as a model system, we demonstrate that incorporating experimentally derived secondary-structure propensities is sufficient to recover accurate interdomain contact maps in a mixed folded– disordered protein.

J-domain chaperones contain a typical, low complexity, glycine and phenylalanine-rich (GF) linker that connects their J-domain with the C-terminal, substrate binding domain and is crucial for cell viability.^7^ In DNAJB6, a highly potent aggregation inhibitor^8^, part of the GF forms a stable α-helix (helix 5 or α5) that blocks the binding of Hsp70 to the J-domain creating a closed/autoinhibited state.^9^ Even though a number of publications investigating the closed/autoinhibited state of DNAJBs are now available,^10,11^ the characterization of the open/uninhibited state in which the GF-linker could serve as a pseudo-substrate for Hsp70^11^, is lacking. Previously, using a construct of the J-domain followed by GF in the absence of helix 5 (JD-GF) as a proxy for the open state we showed by SAXS and NMR spectroscopy that GF partially collapses against JD.^11^ Even though nominally ‘open’, the JD-GF construct shows an order of magnitude decrease in affinity for Hsp70 in comparison to JD alone, presumably due to GF-JD interactions which remain elusive in the atomic level.

In the present study we use solution NMR and CG-simulations to investigate the conformational ensemble sampled by the highly flexible JD-GF construct. Using NMR relaxation methods, we show that specific hydrophobic residues in the otherwise largely disordered GF show reduced motions in the ns timescale. To visualize the interactions between JD and GF we performed CG simulations that reproduce the overall compaction of the molecule but fail to capture the exact interdomain contacts between JD and GF. To overcome this limitation, we introduce backbone dihedral potentials that encode residue-specific secondary-structure propensities derived from NMR chemical shifts into minimal CG simulations resulting in good agreement between the simulation derived JD – GF interdomain contacts and the NMR data. Using this combined NMR and MD approach we generate a physically relevant ensemble for a disordered region in which specific phenylalanine residues in GF form extensive contacts with the J-domain resulting in partially closed states that interfere with Hsp70 binding. More generally, we show that incorporating experimentally-derived local backbone propensities provides an avenue to improving CG descriptions of disordered proteins, enabling access to atomic-level insights that are otherwise difficult to obtain.

## Materials and Methods

### Protein expression and purification

DNAJB6 JD-GF, JD-GF-α5 and DNAJB1 JD-GF constructs were expressed and purified as described previously.^9,11^

### Residual dipolar coupling (RDC) measurements

RDC measurements were collected at 600 MHz on a sample of 200 μM ^13^C, ^15^N-labelled DNAJB6 JD-GF in 20 mM sodium phosphate pH 7.4, 100 mM NaCl, aligned in 14 mg/mL bacteriophage Pf1 (ASLA Scientific) Backbone amide ^1^*D*_NH_ RDCs were measured using the ARTSY pulse sequence.^12^

### Spectral density mapping

Reduced spectral density mapping was performed as in^13^ (Supplementary methods). For well-folded residues the local rotational correlation time that contains contributions from diffusional anisotropy can be obtained from the values of *J*(0) and *J*(*ω*_N_) as:

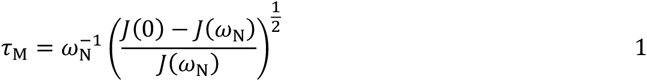

With the value of *τ*_m_ determined, the amplitude of intramolecular motion of the N-H vector can be estimated by^14^:

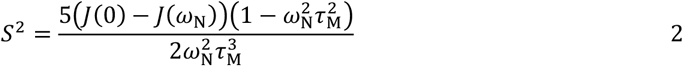

For GF residues the value of *τ*_m_ was fixed to the average value obtained for residues in JD. Given a well defined *τ*_m_ value for folded residues, equation (2) can provide a good approximation of the generalized order parameter square (S^2^) for internal motions in the ns timescale. For example, S^2^ values obtained by equation (2) at one magnetic field show an excellent correspondence with the ones obtained by extensive spectral density mapping at five magnetic fields for the mixed folded protein GCN4^15,16^ which shares a lot of structural and dynamic features with JD-GF.

### Model-free analysis

Model-free analysis was performed in Model-free 4.^17^ Since the closed stated in the JD-GF-α5 construct can be considered as a globular protein with a dynamic linker region (Figure 2A) a global correlation time (*τ*_c_) was used for all residues. For the open JD-GF a global *τ*_c_ was used for JD residues. For GF residues *τ*_c_ was fixed to the value obtained for JD. For residues in the J-domain an axially symmetric diffusion tensor was used to account for the slight anisotropy of the J-domain. The relaxation data were fit to the following models:

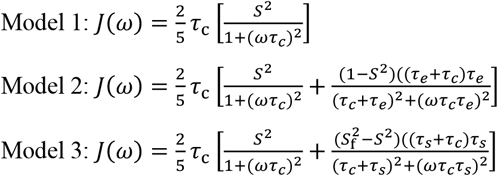

Most of the data for JD and α5 residues can be fitted to the simplest model 1 with only a few residues located in less structured loops showing significant fast motions and thus requiring the use model 2. All GF residues in JD-GF could only be fitted using the extended model-free formalism (model 3). No models that contain exchange contributions to relaxation (*R*_ex_) were used as there is no evidence of such motions in JD-GF. Model selection was performed using the Bayesian Information Criterion (BIC) and errors on the fitted parameters were obtained from 500 Monte Carlo simulations.

### Coarse-grained simulations

CG simulations where each residue was represented as a single bead were performed within the CALVADOS framework^5^ that uses OpenMM as its underlying engine. To allow backbone dihedral angles to be included, the HPS-SS model of Rizuan *et al*.^18^ (Supplementary methods) was implemented as a C++ plugin for OpenMM and is freely available at https://github.com/karamanoslab/OpenMMDihedralPlugin. *ε*_d_ represents the residue-specific parameter that determines the probability of each residue to be in a helical or extended conformation (see Supplementary methods). *ε*_d_ values were modified from the defaults to maximize the agreement with the observed secondary structure propensities calculated by Talos based on the backbone Cα, Cβ, Co, N, HN chemical shifts of JD-GF. The optimized *ε*_d_ values are shown in Supplementary Table 1, with a negative/positive value favoring α-helical/β-strand conformation respectively. *ε*_d_ values are treated here as parameters directly derived from experimentally determined chemical shifts and are unique for the system under study. The dihedral angle potential has little effect on the overall compaction of the ensemble as observed earlier but is effectively modulating the stickiness of each residue/bead by making them more, or less available for interactions. Since the original HPS-SS model is parametrized for Cα dihedral angles we chose to run the CALVADOS simulations with beads centered on the Cα atom of each residue. Backbone dihedral angles where only applied to the GF linker (residues 75 – 98) while the JD was restrained as a rigid body using harmonic restraints. The N-terminal part of helix 3 has a slight kink picked up by its backbone chemical shifts but is fully helical in terms of its Ca configuration. The small helix formed by residues K70 – N74 is too short to be picked up by Talos but has short-range ^1^H-^1^H NOEs fully consistent with a helical conformation. A double well angle potential that describes the angle *θ* between three consecutive residues was also included as described in references [18,19]. When the dihedral angle potential was used the timestep was reduced to 0.005 ps and the number of steps was increased to 2000000 to allow direct comparison with the default CALVADOS simulation (0.01 ps and 1000000 steps respectively). Default values for the stickiness parameter (λ) were used as described in references [4,6].

### Analysis of trajectories

Assignment of helical and extended residues in the simulations was performed as follows: if the dihedral angle φ between 4 consecutive beads (*i, i*+1, *i*+2, *i*+3) falls within the helical (0.1 < φ < 1.5 rad) or extended (-2.5 < φ < -0.5 rad) well, then residues *i*+1, *i*+2 or *i*+1were assigned as pseudo-helical, pseudo-extended respectively. If three consecutive residues in a moving tripeptide window were found to be pseudo-helical/pseudo-extended, the middle residue was assigned as helical/extended. Contact order that emphasises long range interactions was calculated as 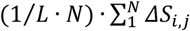 where *N* is the number of contacts for each *i, j* residue pair, Δ*S*_*i, j*_ is the sequence separation and *L* is the total number of residues. Principal component analysis and projection of the trajectory on the principal component vectors was performed in MDanalysis.^20^

## Results and Discussion

### Linker dynamics by NMR relaxation

The dynamics of the DNAJB6 JD-GF were probed by *R*_1_, *R*_2_ and NOE relaxation data at two fields and analysed by spectral density mapping.^13^ The low complexity sequence combined with the intrinsically disordered nature of GF results in highly overlapped resonances for GF residues and therefore the relaxation data were recorded with a triple resonance HNCO readout. Raw relaxation rates shown in Figures 1A – 1C are consistent with a highly ordered JD and a dynamic GF although certain residues in the later seem to be more ordered than others. To gain more insights into the complex dynamics of the GF region the relaxation data were analysed by spectral density mapping^13^ in order to extract the values of *J*(0), *J*(ω_N_) and *J*(ω_H_) (see Methods) which are shown in Figures 1D – 1F. Large *J*(0) and small *J*(ω_H_) values consistent with a stably folded domain are observed for JD residues as expected. Interestingly, residues K70 – N74 in GF also exhibit the same behaviour of those in JD suggestive of a high degree of order for the N-terminus of GF. The patterns of *J*(0) and *J*(ω_H_) are reversed for the rest of GF with small and large values observed respectively in agreement with a highly dynamic region (Figures 1D – 1F). *J*(ω_N_) values for residues D85 - N93 display a rise and fall in amplitude indicative of a sequence dependent variation in nanosecond dynamics (∼1/ω_N_). *J*(ω_H_) values which report on faster, picosecond dynamics (∼1/ω_H_) and are not contaminated by contributions from the global tumbling of the molecule are lower for residues F87, E88, F89, F91, F93 and R94 especially at 600 MHz indicating reduced motions of these residues in the picosecond timescale. Since *ω*_H_*τ*_m_>>1 for JD-GF, *J*(ω_H_) values are mainly affected by internal motions and strongly depend on the order parameter squared (S^2^) which can be estimated purely from *J*(0) and *J*(ω_N_) as shown in Figure 1G. Indeed, S^2^ values for F87, E88, F89, F91, F93 are all above 0.2 consistent with a smaller amplitude of motion for these largely hydrophobic residues.

**Figure 1:**
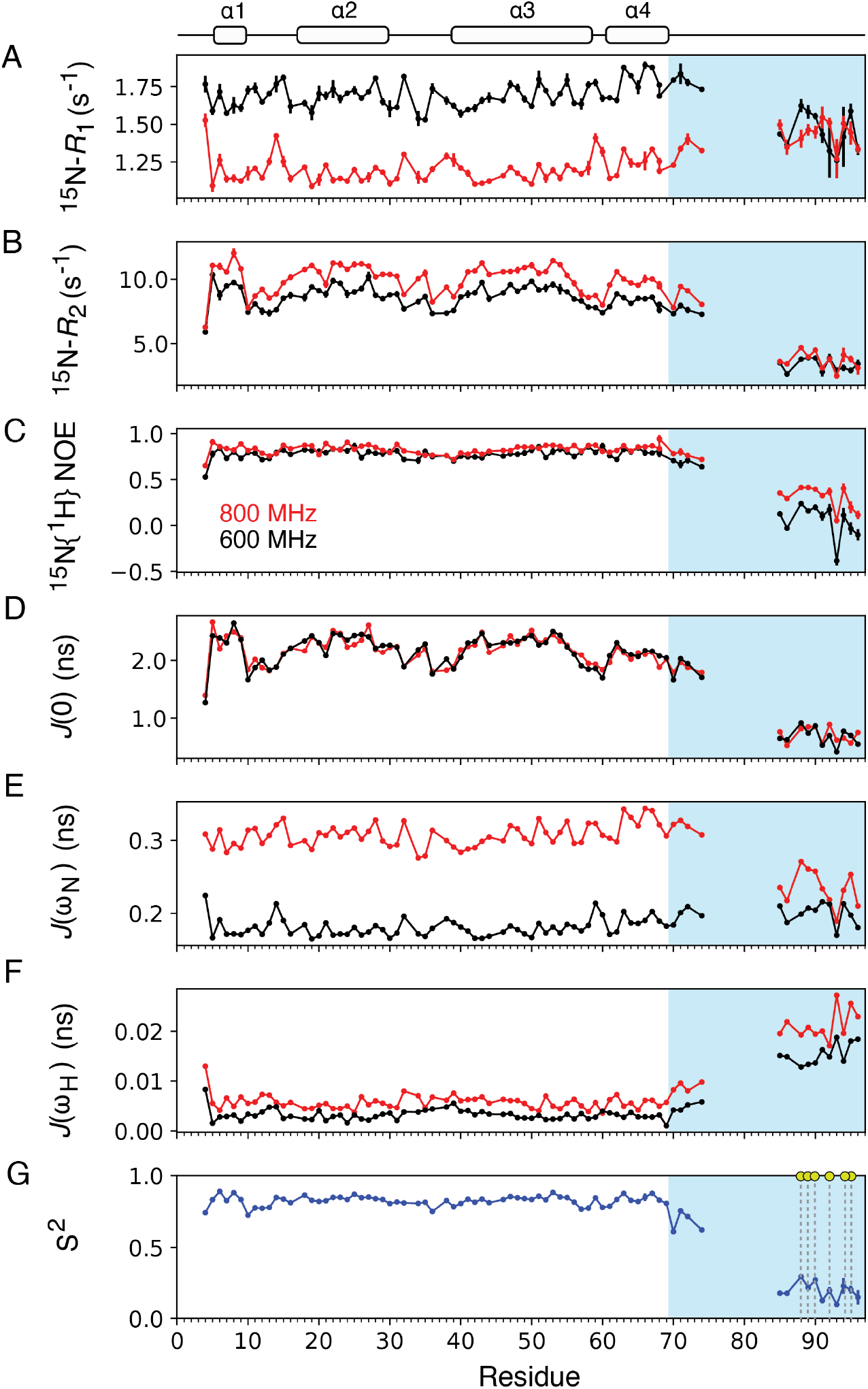
Backbone dynamics of the open JD-GF state. ^15^N *R*_1_ (A), *R*_2_ (B) and ^15^N{^1^H} NOE data (C) collected at 600 (black dots) or 800 (red dots) MHz. *J*(0) (D), *J*(ω_N_) (E) and *J*(ω_H_) (F) values are derived from reduced spectra density mapping. Calculated order parameter squared S^2^ from *J*(0) and *J*(ω_N_) at 800 MHz (eq. 2) are shown in G. GF residues are shown as a cyan box and residues with S^2^ > 0.3 are highlighted with dashed lines/yellow dots.

To gain a more comprehensive understanding of the complex dynamics of the GF region, relaxation measurements at multiple magnetic fields can be performed and provided that the measured spectral densities are linear functions of *ω*, the amplitudes and timescales of motions at various timescales can be extracted with high precision. ^16,21^ Alternatively, a more complicated model-free analysis could be performed, although this is conceptually challenging for fully disordered regions due to the absence of a single tumbling time (*τ*_c_). However, for a system like JD-GF where a relatively short IDR is tethered to and makes contacts with a well-folded domain,^11^ model-free derived motional parameters have been shown to correlate well with those from spectral density mapping at multiple fields.^16^ Based on previous small angle X-ray scattering (SAXS) and chemical shift perturbation (CSP) data^11^ it is safe to assume that GF and JD tumble with the same global correlation time (*τ*_c_) and thus, model-free analysis^22^ can be performed to extract the effective timescales of the internal motions. For helices 1 to 4 a simple model that takes into account the slight anisotropy of the JD diffusion tensor^23^ and only includes a single ordered parameter squared (S^2^) can adequately describe the relaxation data. An effective correlation time for the internal motions (*τ*_e_) in the order of 50 – 100 ps is only needed for loop residues in JD (Figure 1). For GF residues however, a more complicated extended model-free formalism that includes a slow 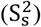 and a fast 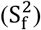 order parameter is required.^14^ For residues K70 – N74 in the GF both 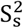 and 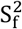 are >0.85 in the open state suggesting a remarkable rigidity of the N-terminal part of the GF-linker. These residues also show ^1^H-^1^H NOEs in the open state that are consistent with a helical structure as seen in various DNAJs (Figure S2A). Moving towards the C-terminal part of GF, S^2^ values drop drastically as expected for an IDR, except for residues F87, E88 (S^2^ ∼ 0.50) and to a smaller extent F89, F91 and R94 (S^2^ ∼ 0.35) which show reduced motions in comparison to other GF residues. A good correlation is observed between the model-free and spectral density mapping derived S^2^ values with a Pearson correlation coefficient of 0.98 and a root-mean-square deviation of 0.06 (Figure S1F). The average effective correlation time for the internal GF motions is on the order of 1 ns, significantly slower than the one observed for the well folded J-domain (*τ*_e_ ∼ 100 ps). Given the complex dynamics of GF, the obtained parameters are not a literal decomposition of their motions, but can be interpreted as transient, large-scale interactions which can obviously take place at various timescales but are captured by a single effective correlation time. Further support that the GF-linker does not adopt a fully random-coil structure stems for ^1^*D*_NH_ residual dipolar coupling (RDC) data (Figure S2B) that show non-zero values for GF residues.

It is instructive to compare the obtained motional parameters for GF residues in the context of the closed state with α5 present (JD-GF-α5 construct, Figure S1). This construct can be considered as a globular protein with a large, disordered linker (Figure 2A) and thus a global correlation time for all residues is appropriate and does not need to be assumed. As expected, the J-domain in the context of JD-GF-α5 behaves similarly to JD-GF (Figures S1 and S3) with helix 5 residues also showing reduced motions consistent with a highly ordered region. GF residues still require an extended model-free formalism to explain their dynamics and show a characteristic drop-off and rise in their S^2^ values (Figure S3F) with a correlation time for the internal motions of around 1.2 ns. Notably, residues K70 – N74 are significantly more dynamic in the closed state showing an 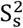 of 0.48-0.75. Taken together, analysis of the NMR relaxation data suggests that hydrophobic GF residues show reduced dynamics in JD-GF, presumably due to interactions with the J-domain.

**Figure 2:**
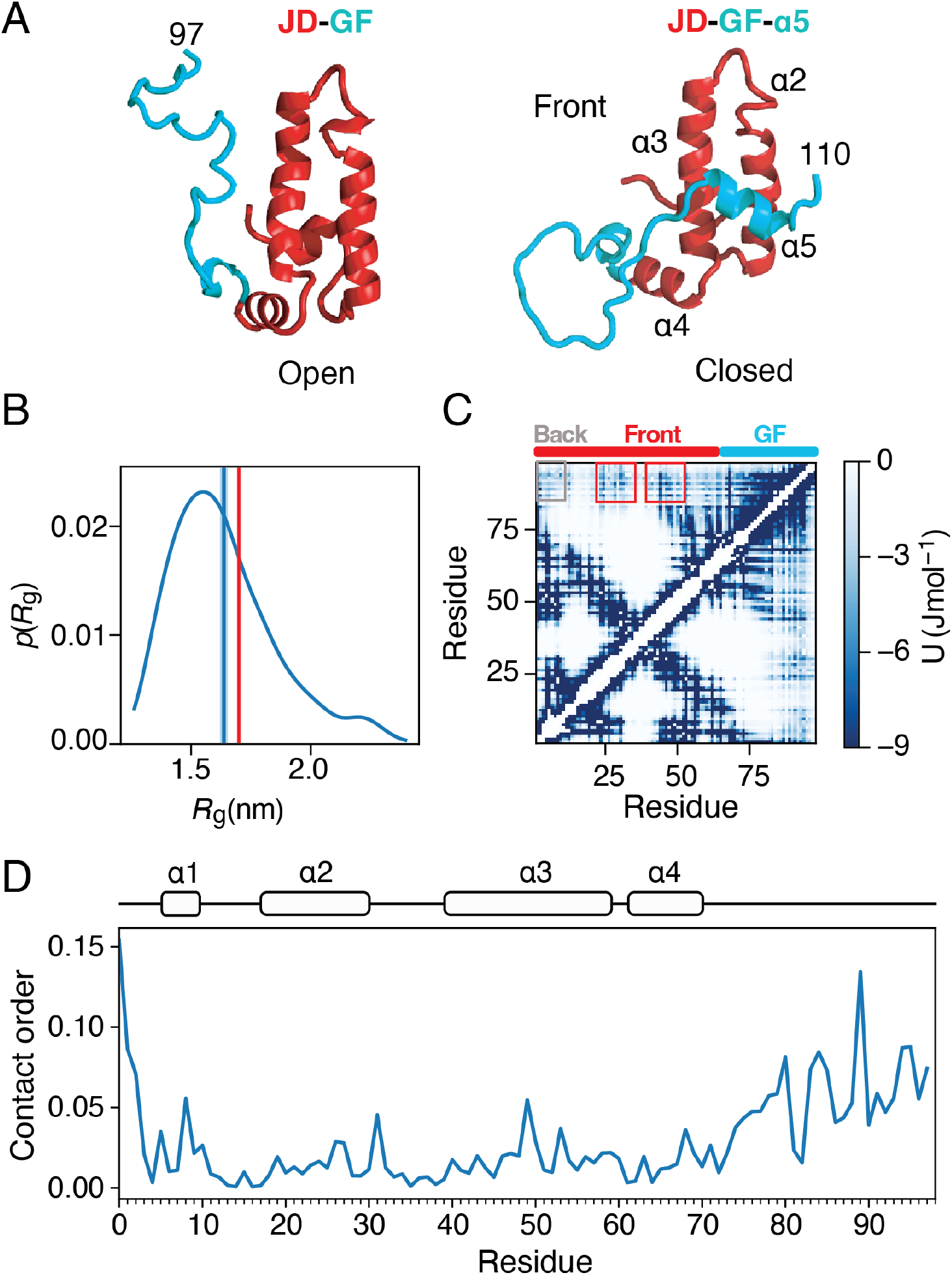
Traditional CG-simulations do not reproduce long-range interactions. (A) Structures of the JD-GF (left) and JD-GF-α5 (right) (JD in red, GF in cyan). (B) Distribution of the radius of gyration from the simulations with the average value and the SAXS determined value indicated as a cyan/red vertical line respectively. (C) Energy maps of the Ashbaugh-Hatch term. Grey/red boxes highlight the back (residues 1-15)/or the front (residues 25-30 and 45-50) of the JD. (D) Contact order values that emphasize long-range interactions (see Methods).

### CG-simulations do not capture the details of the interaction between JD and GF

To structurally visualize the large-scale motions of GF we were interested in generating a conformational ensemble that captures the key properties of the mixed-folded JD-GF. In the presence of experimental data that report on long-range interactions, such as RDCs and paramagnetic relaxation enhancement (PRE) data, such ensembles have been traditionally constructed by fitting the experimental data using an ensemble approach.^24^ However, we specifically avoided to collect PRE data on JD-GF since the highly hydrophobic PRE tags would likely interfere with the hydrophobic interactions that seem most likely to modulate GF motions. Moreover, interproton NOEs require a relatively stable interaction and therefore were only observed for N-but not the C-terminus of GF (Figure S2A). Inspired by the success of the CALVADOS force field^5^ we decided to take a similar, CG-based approach by fine-tuning certain force field parameters to match the available NMR data. Although full-atom MD simulations are possible for systems of this size, they still require significant resources in order to archive the ms timescales relevant for IDRs. On the other hand, CALVADOS uses a single bead representation for each residue that allows sampling of the conformational ensemble of systems that vary in size in a straightforward manner. The NMR relaxation data in Figure 1 suggested that residues K70 - N74 are as rigid as those in the JD and therefore, these were fixed to their native conformations in the simulations. CALVADOS captures the overall compaction of JD-GF, producing an average R_g_ value of 1.65 nm in close agreement with the SAXS determined value of 1.73 nm (Figures 2A, 2B) without the need of any reweighting. However, GF residues transiently interact with the front face of the J-domain that partially exposes various hydrophobic side-chains as evidenced by small chemical shift perturbations in this part of the protein (Figure S4). As seen in Figures 2C and 2D CALVADOS cannot capture these interactions effectively as it predicts significant contacts of the GF with itself and also the back face of the J-domain. Overall, although the global properties of the JD-GF construct are accurately described by the simulations (ensemble compaction measured by *R*_g_), interdomain contacts are not captured.

### Expanding CG-simulations to capture local structure

The inability of the CALVADOS forcefield to generate accurate interdomain contacts is perhaps not surprising as it was only trained using experimental SAXS and PRE data that only report on the global compaction of the system.^5^ We hypothesized that including new terms in the CG simulation that describe local backbone properties would allow us to generate a more accurate ensemble. The backbone chemical shifts of the GF-linker in the open state are not fully random coil but point towards a significant propensity for more extended, β-strand-like conformations.^11^ CALVADOS simulations fail to capture these local structural propensities since the underlying energy function does not contain any terms that bias the backbone towards specific conformations (Figure S4). Capturing secondary structure propensities in single-bead CG-simulations is not straightforward as multiple beads are normally required in order to calculate backbone dihedral angles. Efforts to overcome this limitation of single bead CG-simulations have been presented before by Rizuan et al.^18^ who have used a double well energy potential that allows transition between an α-helical energy well to that of an extended conformation by tuning a residue specific parameter (*ε*_d_). If *ε*_d_ is set to match the known propensities for secondary structure of the various amino acids, this approach can capture the α-helical propensities for a number of IDPs.^18^ However, it is often the case that secondary structure propensities in IDRs cannot be accounted for just purely by their sequence, but are a consequence of the structural intricacies of the system as is the case for JD-GF. Therefore, we decided to tune *ε*_d_ values to match the NMR-determined secondary structure propensity of the GF-linker (Table S1) calculated from its previously determined backbone chemical shifts. In practice this approach could be useful for the plethora of IDRs that have their backbone chemical shifts determined by NMR. Including the backbone dihedral angle term with the optimized εd values in CALVADOS forces GF to populate an extended conformation in its C-terminus, in good agreement with the experimental data (Figure 3A). When compared to the ensemble that was generated without the use of the backbone dihedral angle potential the calculated R_g_ is not significantly smaller (1.63 versus 1.65 nm, 2% difference) (Figures 2B, 3B) showing that the backbone dihedral term does not significantly impact the global compaction of the ensemble. By comparing the chemical shifts of JD alone versus those of JD-GF we have previously shown that the GF-linker locally interacts with the J-domain mainly with the nearby helix 4.^11^ These short-range interactions are also captured by the new simulation that shows good agreement with the experimentally determined chemical shift perturbations (Figure 3C). Overall, the simulations can now capture both the global compaction, the local secondary structure propensities and short-range interactions of GF.

**Figure 3:**
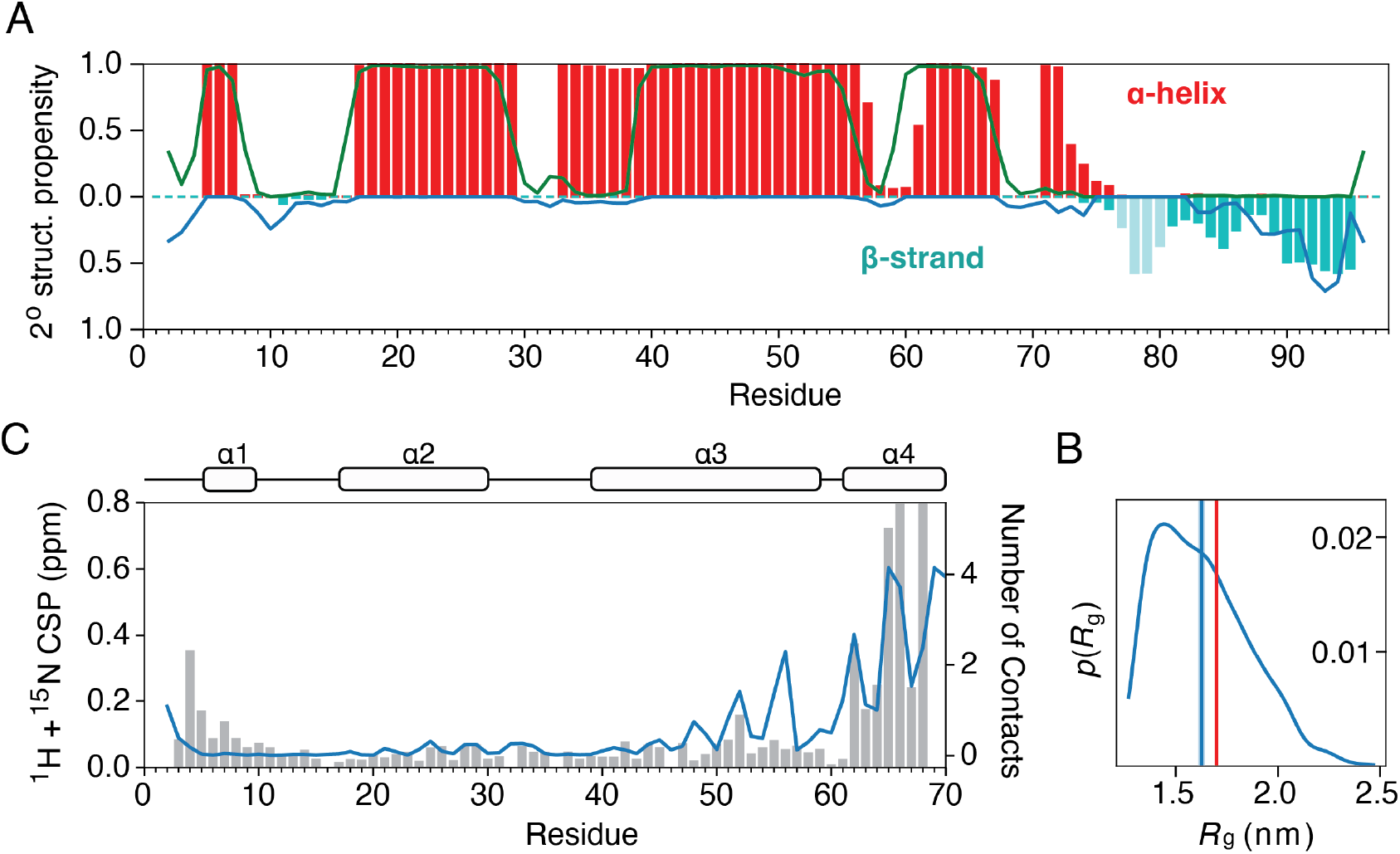
Agreement of the enhanced simulations with experimental data. (A) Secondary structure propensities of JD-GF calculated in the simulation are shown as bars (α-helix, red/β-strand, cyan). Talos-derived secondary structure based on the JD-GF backbone shifts are shown as lines (α-helix, green/β-strand, blue). The JD was restrained as a rigid body and backbone dihedral angles where only applied to the GF linker (residues 75 – 98) (see Materials and methods). Experimental data for residues 75 - 80 are missing due to signal overlap. (B) Distribution of the radius of gyration from the simulations with the average value and the SAXS determined value indicated as a cyan/red vertical line respectively. (C) Combined ^1^H, ^15^N chemical shift perturbations for J-domain resonances in a JD alone versus a JD-GF construct. The number of contacts (<10 Å) each JD residue makes with the GF in the simulation is shown as a cyan line.

### Hydrophobic GF residues lead to partially closed states

We then asked whether the new, enhanced simulations can perform better in terms of generating more accurate long-range interdomain contacts between JD and GF. As seen from the energy maps of Figure 4A when the dihedral angle potential is included the main interdomain interactions involve exposed hydrophobic residues facing the front of the J-domain (helices 2 and 3) in good agreement with small but significant chemical shift perturbations in the same region caused by GF (Figures 4B, S5). Thus, the dihedral angle potential is able to define the correct interdomain contacts by modulating the local conformation of GF residues. Indeed, in the absence of the dihedral potential, GF partially collapses on itself and can form interactions with the nearby regions of the J-domain (Figure 2C). However, when the backbone is forced to adopt an extended conformation in order to satisfy the new dihedral angle potential, the GF can now reach further onto the front of the J-domain. Principal component analysis shows that the compaction of GF onto the front of helices 2 and 3 is the main component of the trajectory, accounting for ∼ 37% of the total motion (Figures 4C, 4D). These motions of the GF relative to the J-domain create states that largely resemble the closed/autoinhibited state even under nominally open conditions, in the absence of helix 5 (Figure 2A) rationalizing the reduced affinity for Hsc70 of the open JD-GF (∼ 400 µM) in comparison to JD alone (∼3 μM).^11^ Analysis of the contact order in the simulations that can emphasize long-range versus short-range interactions, shows spikes for various residues in helices 2 and 3 that all have their side-chains facing the front of the J-domain (Figure 4E). Increased contact order values are also observed for several GF residues, most of which coincide with residues that show reduced motions in the ps – ns timescales as shown by spectral density mapping of NMR data in Figure 1 (Figure 3E). It is thus intriguing to attribute the reduced motions of F87, E88, F89, F91, F93 and R94 to long-range interactions with the J-domain as shown by the CG-simulations. One of the most unexpected results of the analysis of the relaxation data of Figure 1 was the apparent rigidification of residues K70 – N74 in the open state in comparison to the closed one. To investigate the functional role of this collapse we removed the harmonic restraints for these residues to allow them to move freely as the rest of the GF. These new simulations show a significant reduction of the interactions between the linker and the front face of the JD (Figure S6A) suggesting that the conformation of the N-terminal part of the linker is important in positioning the GF ensemble to the front of the J-domain. This process places the hydrophobic GF residues in a position to interact with the exposed hydrophobic residues in JD and could be important in reestablishing autoinhibition after it gets released.

**Figure 4:**
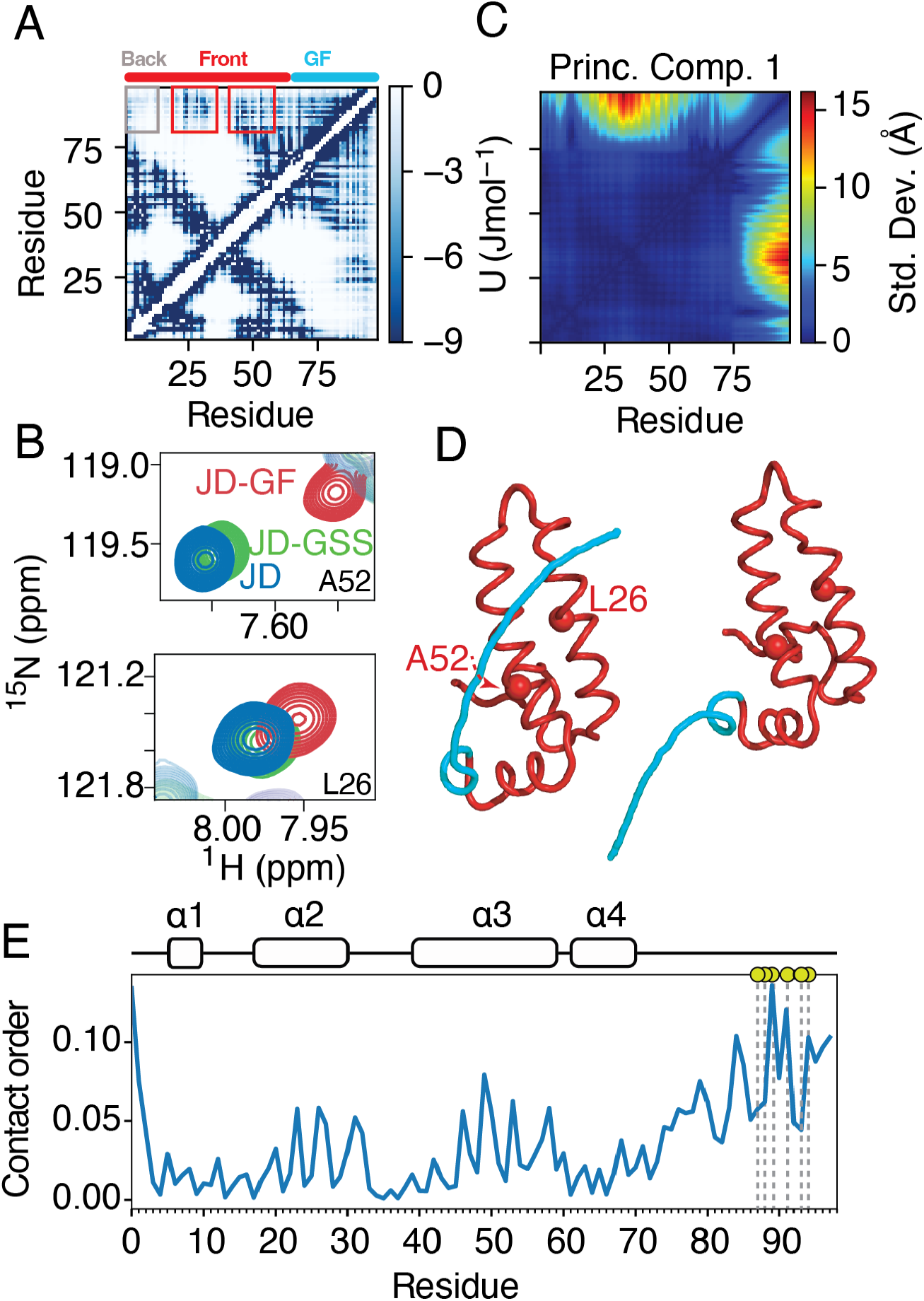
Analysis of the conformational ensemble with the dihedral term included. (A) Energy maps of the Ashbaugh-Hatch term. Grey/red boxes highlight the back (residues 1-15)/or the front (residues 25 -30) of the JD. (B) Regions of the ^1^H-^15^N of JD-GF (red), JD alone (blue) or JD-GSS (green, negative control). (C) Principal component analysis of the resulting trajectory. The principal component vectors are shown as a contact map depicting the change of each distance in relation to all other distances. Principal component 1 describes the compaction of the GF-linker onto the front face of the JD in the open state as shown in (D). L26 and A52 are shown as red spheres. (E) Contact order values that emphasize long-range interactions (see Materials and methods). Yellow spheres and dashed lines represent the GF residues with reduced motions identified by spectral density mapping of the NMR relaxation data (Figure 1).

In order to validate the role of hydrophobic residues in the long-range interdomain interactions, instead of performing multiple single point mutations on JD-GF we decided to swap GF with a fully disordered GSS linker of the same length (JD-GSS construct) as a negative control. As seen in Figure S6B JD-GSS shows an almost complete loss of long-range interdomain interactions in the CG - simulations. This is precisely what we observe experimentally by NMR where the chemical shift perturbations caused on JD resonances by interdomain interactions with GF have almost entirely disappeared in the presence of GSS (Figures S5A, 4B). To further validate the approach, we tested the equivalent JD-GF construct of another class-B DNAJ chaperone, DNAJB1 for which NMR data were available in our lab suggesting that the GF of DNAJB1 is less sticky than that of DNAJB6.^11^ Indeed, the chemical shift perturbations caused by GF on JD are smaller in DNAJB1 JD-GF in comparison to that of DNAJB6 (Figure S7A) showing that long-range interactions are present but less prominent in DNAJB1. CG-simulations portray a similar picture where JD – GF interactions are detected (Figure S7B) but less frequently (Figure S7C).

## Conclusions

Here, we present an approach that enables visualization of accurate interdomain contacts by integrating readily available NMR data that report on local structural features into minimal CG simulations. Guided by NMR data, the approach is used to visualize the interdomain contacts in the JD-GF construct that lead to a significant decrease in affinity for Hsc70. Analysis of NMR relaxation data revealed reduced dynamics for specific hydrophobic residues at both the N- and C-termini of the disordered GF linker, indicating sequence-dependent interactions with the folded J-domain. By tuning residue-specific backbone dihedral parameters (*ε*_d_) to match secondary-structure propensities derived from NMR chemical shifts, accurate interdomain interaction maps between the folded domain and the low-complexity GF region can be obtained. In the absence of explicit backbone dihedral terms, hydrophobic residues within the IDR preferentially collapse onto themselves or nearby sequence segments, preventing access to their native interaction interfaces on the J-domain. Enforcing experimentally determined backbone propensities promotes more extended conformations, thereby enabling these residues to engage hydrophobic surface patches that are otherwise inaccessible computationally. The resulting conformational ensembles rationalize the linker dynamics observed by NMR relaxation and reveal that the nominally open JD–GF construct samples partially closed states which resemble the autoinhibited conformation of JD-GF-α5.

Since backbone chemical shifts are typically among the first NMR observables obtained for intrinsically disordered regions, this strategy provides an experimentally accessible route to improving CG simulations of disordered proteins and mixed folded–disordered systems. Although we focus here on CALVADOS, the proposed approach can be employed in any CG forcefield and can be adapted to optimize against NMR chemical shifts directly.^25^ Regulatory IDRs in signaling proteins, transcription factors, and phase-separating or amyloid proteins are likely to exhibit similar sensitivity to local backbone properties that would be difficult to capture in full-atom simulations due to their increased size. Thus, improved CG models that can reproduce both global and local structural features of the ensemble may broadly improve the accuracy of predicted interdomain interaction maps.

## Data availability

The dihedral potential is freely available as an OpenMM plugin at https://github.com/karamanoslab/OpenMMDihedralPlugin.

## Acknowledgments

We thank Kresten Lindroff-Larsen and Gulio Tessei for helpful discussions. This work was supported by a Sir Henry Dale Fellowship jointly funded by the Wellcome Trust and the Royal Society (Grant Number 223268/Z/21/Z) to TKK. Access to the 700 MHz spectrometer was provided by the MRC Biomedical NMR Centre at the Francis Crick Institute, which receives core funding from Cancer Research UK Grant FC001029; Medical Research Council Grant FC001029; and Wellcome Trust Grant FC001029.

## References

1. Ibanez de Opakua, A. et al. Molecular interactions of FG nucleoporin repeats at high resolution. Nat. Chem. 14, 1278–1285 (2022).

2. Molliex, A. et al. Phase separation by low complexity domains promotes stress granule assembly and drives pathological fibrillization. Cell 163, 123–33 (2015).

3. Borgia, A. et al. Extreme disorder in an ultrahigh-affinity protein complex. Nature 555, 61–66 (2018).

4. Tesei, G. & Lindorff-Larsen, K. Improved predictions of phase behaviour of intrinsically disordered proteins by tuning the interaction range. Open Res Eur 2, 94 (2022).

5. Tesei, G., Schulze, T.K., Crehuet, R. & Lindorff-Larsen, K. Accurate model of liquid-liquid phase behavior of intrinsically disordered proteins from optimization of single-chain properties. Proc Natl Acad Sci U S A 118 (2021).

6. Cao, F., von Bulow, S., Tesei, G. & Lindorff-Larsen, K. A coarse-grained model for disordered and multi-domain proteins. Protein Sci. 33, e5172 (2024).

7. Kampinga, H.H. & Craig, E.A. The HSP70 chaperone machinery: J proteins as drivers of functional specificity. Nat. Rev. Mol. Cell Biol. 11, 579 EP–592 (2010).

8. Kakkar, V. et al. The S/T-Rich Motif in the DNAJB6 Chaperone delays polyglutamine aggregation and the onset of disease in a mouse model. Mol. Cell 62, 272–283 (2016).

9. Karamanos, T.K., Tugarinov, V. & Clore, G.M. Unraveling the structure and dynamics of the human DNAJB6b chaperone by NMR reveals insights into Hsp40-mediated proteostasis. Proc. Natl. Acad. Sci. U. S. A. 116, 21529–21538 (2019).

10. Abayev-Avraham, M., Salzberg, Y., Gliksberg, D., Oren-Suissa, M. & Rosenzweig, R. DNAJB6 mutants display toxic gain of function through unregulated interaction with Hsp70 chaperones. Nat Commun 14, 7066 (2023).

11. Hobbs, B. et al. A low-complexity linker as a driver of intra- and intermolecular interactions in DNAJB chaperones. Nat Commun 16, 5070 (2025).

12. Fitzkee, N.C. & Bax, A. Facile measurement of 1H-15N residual dipolar couplings in larger perdeuterated proteins. J. Biomol. NMR 48, 65–70 (2010).

13. Farrow, N.A., Zhang, O., Szabo, A., Torchia, D.A. & Kay, L.E. Spectral density function mapping using 15N relaxation data exclusively. J. Biomol. NMR 6, 153–62 (1995).

14. Lefevre, J.F., Dayie, K.T., Peng, J.W. & Wagner, G. Internal mobility in the partially folded DNA binding and dimerization domains of GAL4: NMR analysis of the N-H spectral density functions. Biochemistry 35, 2674–86 (1996).

15. Bracken, C., Carr, P.A., Cavanagh, J. & Palmer, A.G., 3rd. Temperature dependence of intramolecular dynamics of the basic leucine zipper of GCN4: implications for the entropy of association with DNA. J Mol Biol 285, 2133–46 (1999).

16. Gill, M.L., Byrd, R.A. & Palmer, A.G., III. Dynamics of GCN4 facilitate DNA interaction: a model-free analysis of an intrinsically disordered region. Phys. Chem. Chem. Phys. 18, 5839–49 (2016).

17. Mandel, A.M., Akke, M. & Palmer, A.G., 3rd. Backbone dynamics of Escherichia coli ribonuclease HI: correlations with structure and function in an active enzyme. J Mol Biol 246, 144–63 (1995).

18. Rizuan, A., Jovic, N., Phan, T.M., Kim, Y.C. & Mittal, J. Developing bonded potentials for a coarse-grained model of intrinsically disordered proteins. J. Chem. Inf. Model. 62, 4474–4485 (2022).

19. Kim, Y.C. & Hummer, G. Coarse-grained models for simulations of multiprotein complexes: application to ubiquitin binding. J Mol Biol 375, 1416–33 (2008).

20. Michaud-Agrawal, N., Denning, E.J., Woolf, T.B. & Beckstein, O. MDAnalysis: a toolkit for the analysis of molecular dynamics simulations. J. Comput. Chem. 32, 2319–27 (2011).

21. Mayo, K.H., Daragan, V.A., Idiyatullin, D. & Nesmelova, I. Peptide internal motions on nanosecond time scale derived from direct fitting of 13C and 15N NMR spectral density functions. J Magn Reson 146, 188–95 (2000).

22. Lipari, G. & Szabo, A. Model-Free approach to the interpretation of nuclear magnetic-resonance Relaxation in macromolecules.1. Theory and range of validity. J. Am. Chem. Soc. 104, 4546–4559 (1982).

23. Tjandra, N., Feller, S.E., Pastor, R.W. & Bax, A. Rotational diffusion anisotropy of human ubiquitin from N-15 NMR relaxation. J. Am. Chem. Soc. 117, 12562–12566 (1995).

24. Clore, G.M. & Iwahara, J. Theory, practice, and applications of paramagnetic relaxation enhancement for the characterization of transient low-population states of biological macromolecules and their complexes. Chem Rev 109, 4108–39 (2009).

25. Cullen, M. et al. Integrating NMR restraints into coarse-grained simulations: toward accurate conformational ensembles of complex protein systems. bioRxiv, 2025.12.22.695971 (2025).

