## Supplementary Data for "Accurate interdomain contacts in mixed folded proteins from NMR-guided coarse-grained simulations"

**Experimental methods, 1 Supplementary Table and 7 Figures**

**Supplementary methods**

**Spectral density mapping**. The measured $^{15}$N relaxation rates can be described as:

$$R_1 = \left(\frac{d^2}{4}\right)\left(3J(\omega_N) + 6J(\omega_N + \omega_H) + J(\omega_N - \omega_H)\right) + c^2 J(\omega_N) \quad 1$$

$$R_2 = \left(\frac{d^2}{8}\right)\left(4J(0) + 3J(\omega_N) + 3J(\omega_N + \omega_H) + 6J(\omega_H) + J(\omega_N - \omega_H)\right) + \left(\frac{c^2}{6}\right)\left(J(0) + 6J(\omega_N)\right) \quad 2$$

$$\mathrm{NOE} = 1 + \left(\frac{d^2}{4R_1}\right)\left(\frac{\gamma_H}{\gamma_N}\right)\left(J(\omega_H + \omega_N) - J(\omega_H - \omega_N)\right) \quad 3$$

where $d = (\mu_0 h \gamma_H \gamma_N)\langle r_{NH}^{-3}\rangle$, $c = \omega_N \Delta\sigma/\sqrt{3}$, $\mu_0$ is the permeability of the vacuum, $h$ Planck's constant, $\gamma_H$, $\gamma_N$ are the gyromagnetic ratios of $^1$H and $^{15}$N respectively, $r_{NH}$ = 1.02 Å and $\Delta\sigma$ = -172 ppm is the chemical shift anisotropy. Reduced spectral density mapping was performed as in Supplementary reference [[1]] with values of $J(0)$, $J(\omega_N)$ and $J(\omega_H)$ are obtained as:

$$\sigma_{NH} = R_1(\mathrm{NOE} - 1)\,\gamma_N/\gamma_H \quad 4$$

$$J(\omega_H) = \frac{4\sigma_{NH}}{5d^2} \quad 5$$

$$J(\omega_N) = \frac{4R_1 - 5\sigma_{NH}}{3d^2 + 4c^2} \quad 6$$

$$J(0) = \frac{6R_2 - 3R_1 - 2.72\sigma_{NH}}{3d^2 + 4c^2} \quad 7$$

**Coarse-grained simulations.** The HPS-SS potential has the form[2]:

$$U_{dihed}(\varphi, \varepsilon_d) = -\ln\left[U_{dihed,\alpha}(\varphi, \varepsilon_d) + U_{dihed,\beta}(\varphi, \varepsilon_d)\right] \quad 8$$

Where $\varphi$ is the dihedral angle of four consecutive residues and $U_{dihed,\alpha}$ , $U_{dihed,\beta}$ are the functions that represent the helical and extended conformations. $\varepsilon_d$ represents the residue-specific parameter that determines the probability of each residue to be in a helical or extended conformation. The helical part is given by:

$$U_{dihed,\alpha}(\varphi, \varepsilon_d) = e^{-k_{\alpha,1}(\varphi - \varphi_{\alpha,1})^2 - \varepsilon_d} + e^{-k_{\alpha,2}(\varphi - \varphi_{\alpha,2})^4 + \varepsilon_0} + e^{-k_{\alpha,2}(\varphi - \varphi_{\alpha,2} + 2\pi)^4 + \varepsilon_0} \quad 9$$

where $k_{\alpha,1}$ = 11.4 rad$^{-2}$, $k_{\alpha,2}$ = 0.15 rad$^{-4}$, $\varphi_{\alpha,1}$ = 0.90 rad, $\varphi_{\alpha,2}$ = 1.02 rad and $\varepsilon_0$ =0.27. The extended part of the function is:

$$U_{dihed,\beta}(\varphi, \varepsilon_d) = e^{-\mathrm{k}_{\beta,1}(\varphi-\varphi_{\beta,1})^2+\varepsilon_d} + e^{-\mathrm{k}_{\beta,2}(\varphi-\varphi_{\beta,2}-2\pi)^2+\varepsilon_1+\varepsilon_d} + e^{-\mathrm{k}_{\beta,2}(\varphi-\varphi_{\beta,2})^4+\varepsilon_2} + e^{-\mathrm{k}_{\beta,2}(\varphi-\varphi_{\beta,2}-2\pi)^4+\varepsilon_2} \quad 10$$

where $k_{\beta,1}$ = 1.80 rad$^{-2}$, $k_{\beta,2}$ = 0.65 rad$^{-4}$, $\varphi_{\beta,1}$ = -1.55 rad $\varphi_{\beta,2}$ = -2.50, rad $\varepsilon_1$ =0.14 and $\varepsilon_2$ =0.40. $\varepsilon_d$ values were modified from the default values to maximise the agreement with the observed secondary structure propensities calculated by Talos based on the backbone Cα, Cβ, Co, N, HN chemical shifts of JD-GF.

### Supplementary Tables

Table 1: Optimised $\varepsilon_d$ values for the HPS-SS backbone dihedral potential.

| Residue | $\varepsilon_d$ |
|---|---|
| R | 3.00 |
| D | 0.76 |
| N | 1.32 |
| E | -0.58 |
| K | 0.54 |
| H | 0.11 |
| Q | 0.13 |
| S | 0.89 |
| C | 1.73 |
| G | 2.89 |
| T | 1.98 |
| A | -1.65 |
| M | -0.38 |
| Y | 0.94 |
| V | 0.79 |
| W | 0.37 |
| L | -0.90 |
| I | -0.12 |
| P | 2.65 |
| F | 0.94 |

### Supplementary Figures

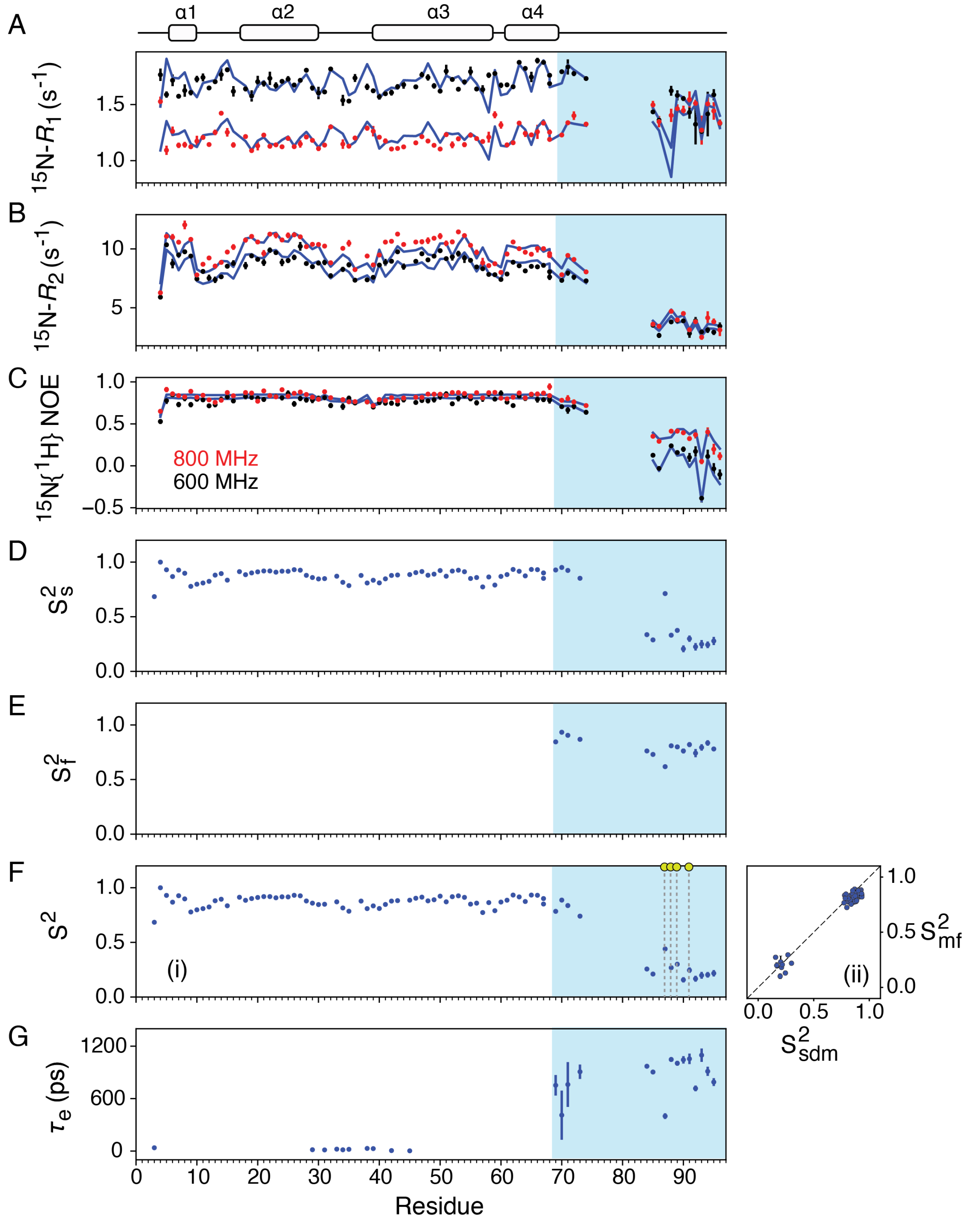


**Figure S1: Model-free analysis of the open JD-GF construct.** $^{15}$N $R_1$ (A), $R_2$ (B) and $^{15}$N{$^1$H} NOE data (C) collected at 600 (black dots) or 800 (red dots) MHz with model-free predicted values shown as blue lines. Calculated order parameter squared for the slow ($S^2_s$), fast ($S^2_f$), the generalised order parameter $S^2 = S^2_s S^2_f$ and the correlation time for the internal motion ($\tau_e$) are shown in D, E, F (panel i) and G respectively. Panel F(ii) shows the corelation between the spectral density derived $S^2$ values with those derived from the model-free analysis (RMSD 0.06, Pearson's corelation coefficient 0.98). An axially symmetric diffusion tensor was used for JD residues and a fully isotropic extended spectral density function for those in GF. When an extended model-free formalism was used $\tau_e$ specifically refers to the slow motion ($\tau_e = \tau_s$).

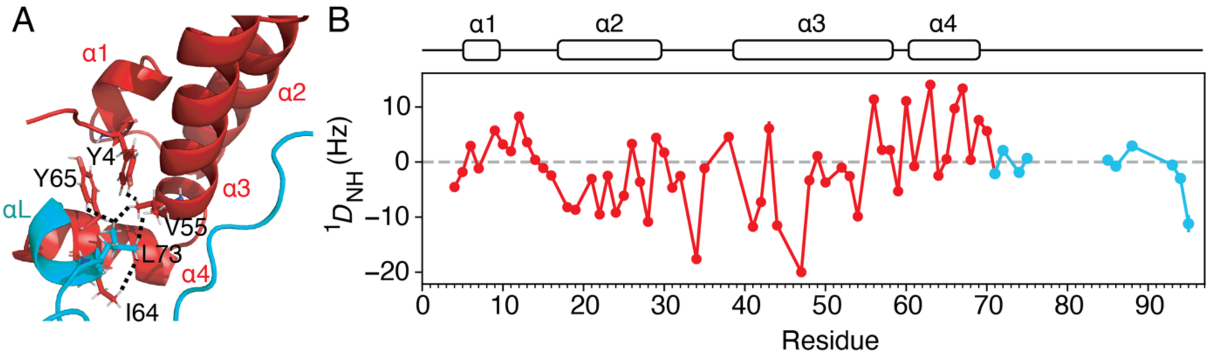


**Figure S2: Structural analysis of the GF region**. A cartoon representation of JD-GF with the J-domain and the GF shown in red and cyan respectively. The observed $^1H$ – $^1H$ NOEs involving residues K70 – N74 (labelled as αL) are consistent with a helical configuration of this segment (labelled as αL) and are shown as black dashed lines.[3] (B) $^1D_{NH}$ RDCs for JD-GF measured in 14 mg/mL Pf1 bacteriophage. Residues in the J-domain are shown in red and those in GF in cyan.

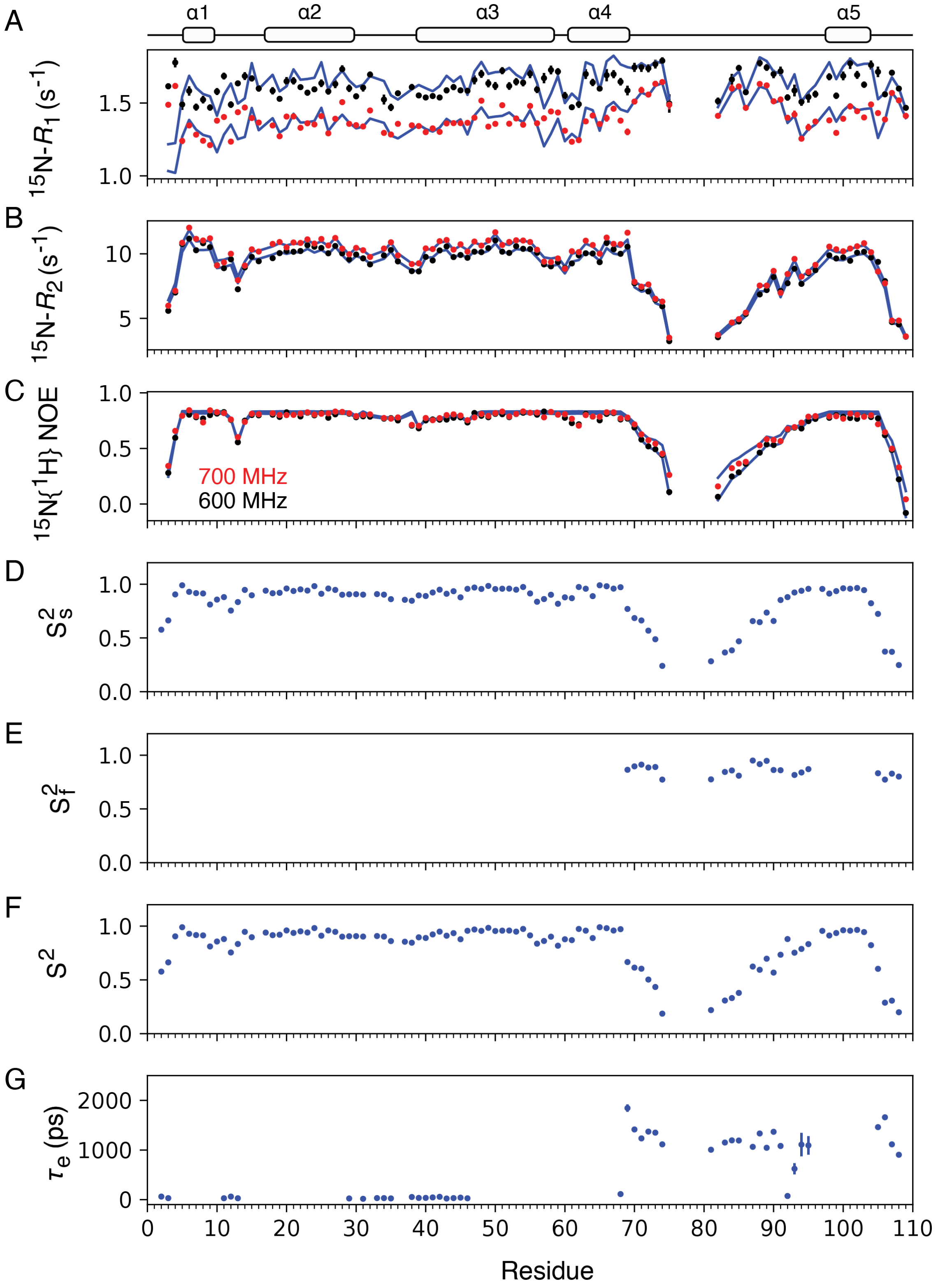


**Figure S3: Model-free analysis of the closed JD-GF-α5 construct.** $^{15}$N $R_1$ (A), $R_2$ (B) and $^{15}$N{$^1$H} NOE data (C) collected at 600 (black dots) or 700 (red dots) MHz with model-free predicted values shown as blue lines. Calculated order parameter squared for the slow ($S_s^2$), fast ($S_f^2$), the generalised order parameter $S^2 = S_s^2 S_f^2$ and the correlation time for the internal motion ($\tau_e$) are shown in D, E, F and G respectively. An axially symmetric diffusion tensor was used for JD residues and a fully isotropic extended spectral density function for those in GF. When an extended model-free formalism was used τe specifically refers to the slow motion ($\tau_e = \tau_s$).

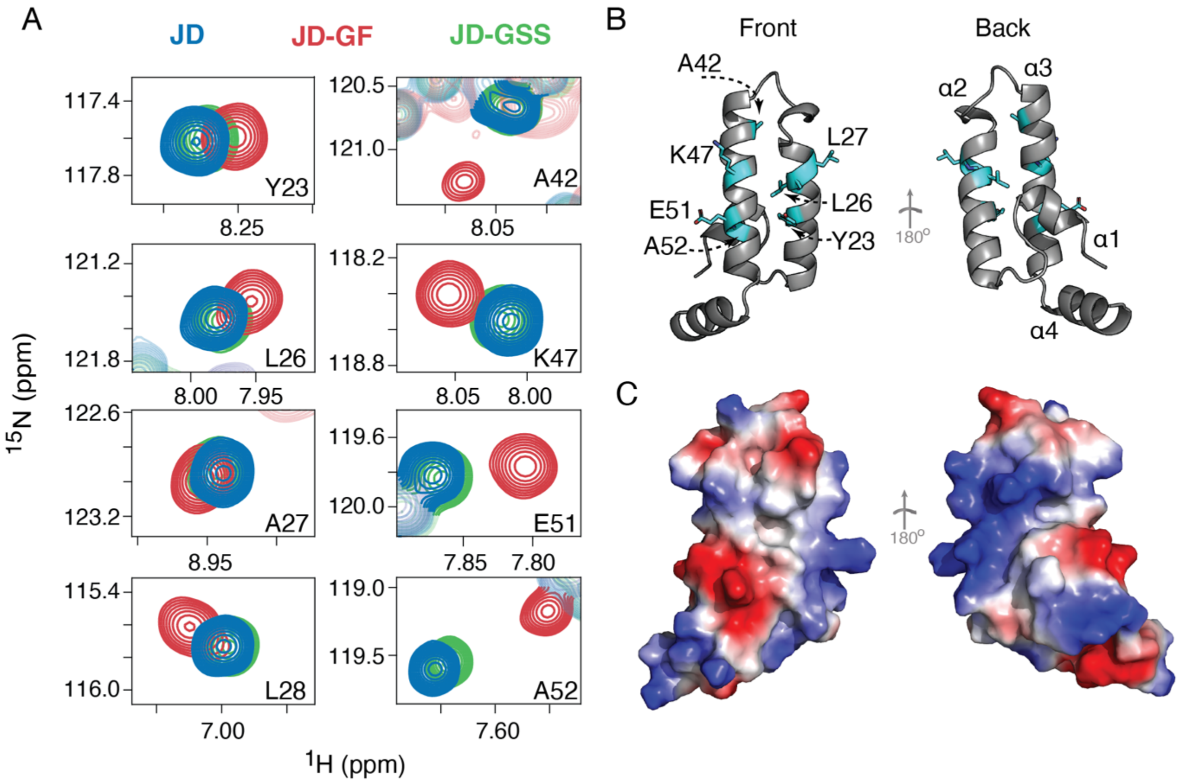


**Figure S4: Contacts between the GF-linker and the hydrophobic front face of the JD.** (A) Regions of the $^{1}H$-$^{15}N$ of JD-GF (red), JD alone (green) or JD-GSS (green) for various JD residues. The CSPs caused by GF essentially disappear in the presence of a disordered Gly-Ser-Ser repeats of the same length in JD-GSS. The residues highlighted in (A) are shown in cyan on the structure of JD in (B). The electrostatic surface potential of JD is shown in (C) using the same front and back poses as in (B) (red – negatively charged, white – hydrophobic, blue – positively charged).

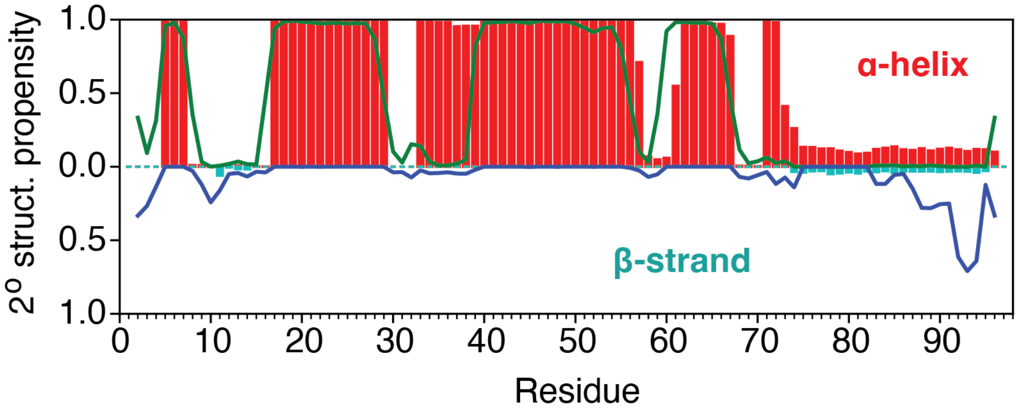


**Figure S5: Conformational analysis of the JD-GF ensemble with no backbone dihedral angles included.** Secondary structure propensities calculated in the CG-simulation are shown as bars (α-helix, red/β-strand, cyan). Talos-derived secondary structure[4] based on the JD-GF backbone shits are shown as lines (α-helix, green/β-strand, blue).

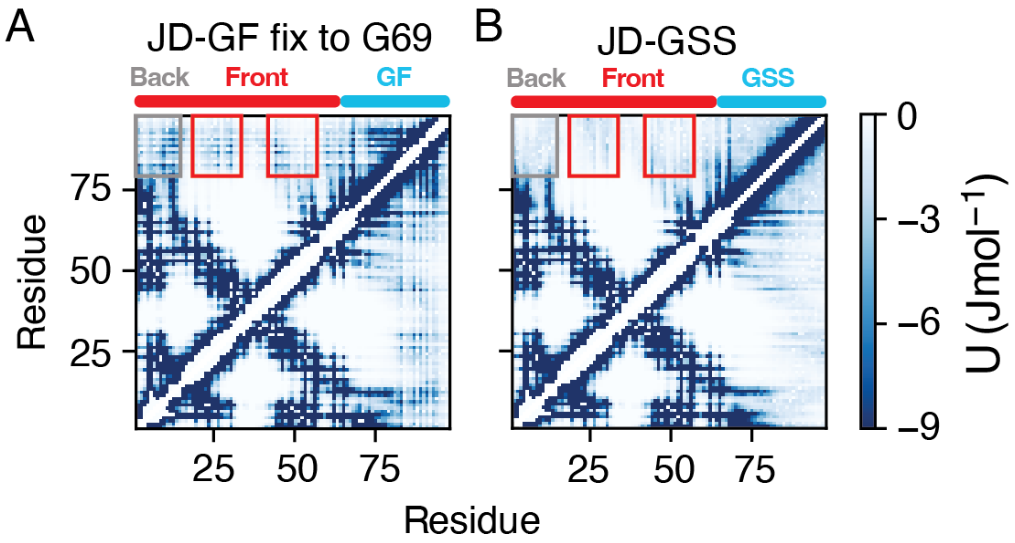


**Figure S6: Modulating interdomain contacts.** Energy maps of the Ashbaugh-Hatch term for a simulation where harmonic restraints were applied only up to residue G69 in JD-GF (A) or for the JD-GSS construct in which the GF is swapped with Gly-Ser-Ser repeats of the same length as GF (B). Grey/red boxes highlight the back (residues 1-15)/or the front (residues 25 - 30) of the JD.

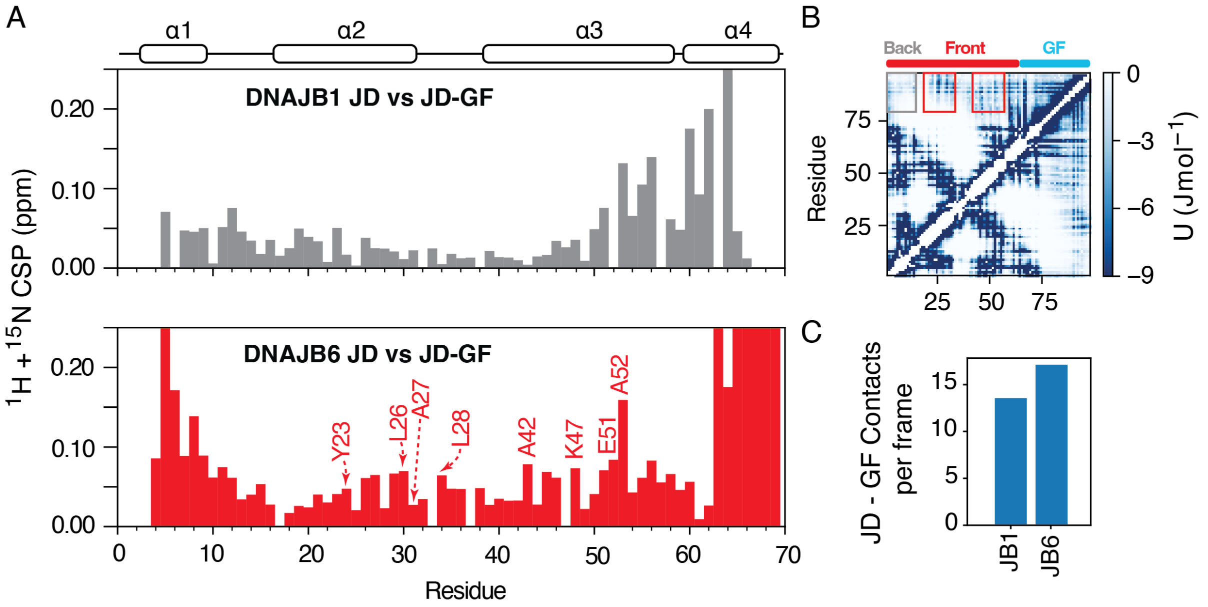


**Figure S7: Interdomain contacts vary between class-B DNAJs.** Combined $^{1}H$, $^{15}N$ chemical shift perturbations for JD resonances in a JD alone versus a JD-GF construct for DNAJB1 (top panel) or DNAJB6(bottom panel). The observed CSPs are generally smaller for DNAJB1 suggesting a smaller number of contacts between JD and GF in comparison to DNAJB6. Residues shown in Figure S5A are highlighted on the bottom graph. (B) Energy map of the Ashbaugh-Hatch term for the DNAJB1 JD-GF simulation. (C) Number of JD – GF contacts per frame for DNAJB1 and DNAJB6.

**Supplementary References**


1. Farrow, N.A., Zhang, O., Szabo, A., Torchia, D.A. & Kay, L.E. Spectral density function mapping using $^{15}N$ relaxation data exclusively. *J. Biomol. NMR* **6**, 153-62 (1995).
2. Rizuan, A., Jovic, N., Phan, T.M., Kim, Y.C. & Mittal, J. Developing bonded potentials for a coarse-grained model of intrinsically disordered proteins. *J. Chem. Inf. Model.* **62**, 4474-4485 (2022).
3. Hobbs, B. et al. A low-complexity linker as a driver of intra- and intermolecular interactions in DNAJB chaperones. *Nat. Commun.* **16**, 5070 (2025).
4. Shen, Y., Delaglio, F., Cornilescu, G. & Bax, A. TALOS+: a hybrid method for predicting protein backbone torsion angles from NMR chemical shifts. *J. Biomol. NMR* **44**, 213-223 (2009).
